# Latent Literacy in Minimally Verbal Autistic Individuals Revealed by Eye Gaze

**DOI:** 10.1101/2025.11.09.687411

**Authors:** Keren Ellert, Hannah S. Sykes-Haas, Niva Shefer-Kaufmann, Yoram S. Bonneh

## Abstract

Minimally verbal autistic individuals (mvASD) are often presumed to have severe cognitive and language impairments based on their poor performance on standardized assessments requiring voluntary motor responses, such as pointing. However, emerging evidence suggests that these individuals may possess latent cognitive abilities. Here, we introduce the Cued Looking Paradigm (CLP), a novel eye-tracking method that bypasses motor requirements by capturing automatic gaze responses to language-based stimuli. In our study, 35 minimally verbal autistic adolescents and adults were presented with spoken or written words, followed by a pair of images (target and foil) as their eye movements were recorded. Among mvASD participants with usable eye-tracking data (n = 30), the majority (80%) demonstrated hidden receptive language and reading abilities, as evidenced by eye-gaze measures, including temporal dynamics and spatial displacement, that were comparable to those observed in neurotypical controls. In contrast, the same mvASD individuals averaged only 57% accuracy when asked to read and point to the target picture, revealing a significant gap between reporting via pointing and actual lexical-semantic knowledge. Furthermore, pupil dilation analysis during tasks indicated reduced arousal recruitment in mvASD participants, potentially implicating dysregulation of the locus coeruleus-norepinephrine (LC-NE) system associated with the performance gap between pointing and eye-gaze. These findings challenge assumptions of global intellectual limitation while confirming specific lexical-semantic competence among mvASD individuals. Results highlight the need for, and provide, alternative- assessments that bypass manual motor responses. The CLP shows promise for revealing cognitive and language abilities, with important implications for both research and education.

**Significance Statement:** Standard language tests implicitly assume that a person can point or speak. Although minimally verbal individuals can point, this response may not reliably reflect comprehension. Using a simple eye-tracking task that replaces pointing with automatic gaze shifts, we show that most mvASD participants accurately match spoken or written words to pictures—even though they fail the same task when pointing is required. This finding challenges the assumption that absence of speech is typically associated with absence of understanding and reveals bias in common assessments. Tools that bypass manual motor demands by using eye movements, such as the Cued Looking Paradigm, together with changes in assessment and intervention, could transform diagnosis, guide education, and open new research avenues on covert language processing.

## Introduction

Minimally verbal autistic individuals (mvASD), estimated about 30% of the autism spectrum (1, 2) have often been grouped together with autistic individuals with cooccurring intellectual disabilities (3). This categorization is primarily based on their poor performance on conventional assessments (4), such as the PPVT-4 or Raven’s Progressive Matrices, which depend on voluntary motor responses (5). While one explanation attributes this to true deficits in cognition and receptive language (6), such an interpretation may overlook the heterogeneity within the mvASD population (3). This heterogeneity is supported by reports from caregivers and clinicians of clear signs of comprehension of spoken or printed words. Indeed, a lack of spoken language and inconsistent behavior do not necessarily indicate impaired comprehension or cognition (7). Supporting this view, previous research (5) found that two-thirds of mvASD children scored within the typical age-range (>70) on a modified version of the Raven’s Colorful Progressive Matrices (RCPM) (3, 8, 9) (see also Sykes-Haas & Bonneh, preprint (10)). These findings suggest that poor performance on standardized tests may not reflect a lack of ability, but rather difficulties in demonstrating cognitive skills via pointing, leading to floor performance and underestimated competence (4, 5). This pattern is echoed in our recent study (10) in which mvASD children (ages 6–11) frequently *fixated* on the target in a simple visual oddball task but failed to *point* to it, underscoring a striking gap between perceptual capacity and voluntary action. These considerations align with broader neurocognitive accounts of language and conceptual processing in neurological populations, which emphasize the need for assessment methods that are sensitive to preserved knowledge despite impaired overt responses (e.g.,(11). Intervention studies further indicate that even mvASD children can show increased social engagement and communicative exchanges when provided with structured, peer-mediated support, revealing abilities that are not apparent in standard testing contexts (12, 13).

We propose that for a subset of mvASD individuals, the primary barrier to communication may be a profound difficulty in executing intentional goal-directed motor actions in the absence of external structure or support. This difficulty may reflect dysregulated action selection, limiting the production of appropriate motor behavior and constituting an “output problem” rather than a deficit in understanding (see (14) for a framework of action selection under competing affordances). Consistent with this view, evidence of atypical motor variability in autism suggests that impaired movement control can mask underlying competence in standard performance-based assessments (15). Importantly, such motor variability is expected to disproportionately impair overt, precision-dependent actions (e.g., pointing or speech), while leaving lower-threshold readouts of action selection—such as gaze allocation—relatively intact. Eye gaze is a highly overlearned and largely automatic behavior, making it a particularly suitable channel for revealing latent literacy when actions such as pointing remain effortful and non-automatized (16). Here, we use the term voluntary to refer to internally initiated, goal-directed actions in the absence of external scaffolding, without implying reduced intent or motivation.

Taken together, the aforementioned findings and observations highlight the need for alternative assessment approaches that among others may rely on involuntary, automatized, or physiological responses. Such methods may bypass impaired voluntary motor control or disrupted affordance competition, and reveal cognitive abilities that remain hidden in tasks requiring voluntary actions, such as pointing (5, 17, 18). To address this need, we introduce the **Cued Looking Paradigm (CLP)**, a novel eye-tracking method designed to uncover receptive language and reading skills in individuals with mvASD. Unlike the Looking-While-Listening (LWL) paradigm (19) which may still involve more controlled search behavior, CLP may reduce task load by presenting the auditory or written cue *before and separately from* the image pair. This design may promote automatic gaze responses without engaging conscious decision-making or deliberate motor planning (16). We hypothesize that due to these aforementioned modifications the CLP will more accurately capture lexical-semantic competence in minimally verbal individuals with autism.

### Evidence for Hidden Cognition and Action-Control Deficits in mvASD

Clinical observations suggest that some preverbal toddlers and older children with mvASD demonstrate rapid eye gaze to correct objects when prompted, even when they fail to point (author NSK, personal clinical experience). Some mvASD individuals can independently communicate through typing (20). Unlike the discredited facilitated communication method (21), these individuals generate meaningful text without physical assistance (20), and some report difficulties initiating and controlling intentional actions (22), such as pointing inaccurately despite knowing the correct response (23). Jaswal et al. (24, 25) provide indirect support for this dissociation. Eye-tracking during letterboard use showed that mvASD participants fixated on each letter just before touching it, with timing like neurotypical spellers. A follow-up study using a fixed iPad ruled out manual prompting by assistant and found independent letter selection and faster responses for meaningful sentences and common letter pairs, consistent with English orthographic knowledge. These findings suggest lexical– semantic literacy in low-verbal autistic individuals. Additionally, some researchers have linked autism to catatonia, a condition where individuals retain awareness but struggle to initiate actions (26). Across these findings, a common theme emerges: a consistent gap between observed overt behavior and latent cognitive abilities.

Despite this compelling evidence for latent cognitive abilities in mvASD, we do not endorse the notion that mvASD individuals are simply typically developing individuals whose intact cognition and perception are merely masked by output barriers. We acknowledge that these individuals face substantial expressive challenges that are not yet fully understood or characterized and could largely vary across individuals. Our position is that certain research approaches or experimental contexts may allow previously unobservable or latent capacities to become evident. The nature and developmental status of these capacities—whether potential, partially or substantially developed, or contextually hidden—remain open empirical questions rather than theoretical assumptions of intact or typical functioning.

### Limitations of ERP and LWL Approaches

Several studies have used ERP methods to explore receptive language in mvASD without requiring active responses. Most focus on the N400 component, which signals mismatches between spoken words and visual stimuli (18, 27, 28). However, results have been mixed; only one study found significant semantic processing in a group of 20 mvASD children (18). Group-level findings suggest some sensitivity to lexical processing, but individual variability and task constraints remain limiting. Eye-tracking offers a noninvasive, behavior-independent alternative for assessing cognition like in the aforementioned LWL paradigm (29). However, Plesa-Skwerer et al. (19) did not find eye-gaze performance more informative than pointing. We suggest that these findings may reflect task complexity of the LWL rather than true ability and that the novel CLP overcomes these experimental challenges.

### The current study

In the present study, 35 mvASD adolescents and adults completed two CLP conditions (spoken-word and written-word trials). In matched conditions with identical cues and response stimuli, participants were also asked to respond by pointing to laminated cards, simulating standard educational assessments (e.g., PPVT-4). This design allowed us to directly compare gaze-based and manual responses within the same individuals. Strikingly, the CLP revealed lexical-semantic knowledge that was not evident from pointing responses, suggesting that gaze-based methods provide a more accurate estimate of cognitive abilities in mvASD.

## Methods

### Participants

Thirty-five mvASD and 30 age-matched TD participants (15–30 years), all from Hebrew-speaking homes, were enrolled (Table 1). mvASD participants had official DSM-IV/5 diagnoses, used <30 functional words by caregiver report (confirmed in a ∼15-min experimenter interaction), and caregivers completed the SCQ; mvASD were recruited by word of mouth via clinics/special-education schools and excluded for epilepsy or ability to read aloud (TD: no neurologic/psychiatric history). The study was approved by the Bar-Ilan University Ethics Committee and written consent was obtained; five mvASD participants were excluded during eye-tracking quality control, leaving 30 mvASD and 30 TD in the final analysis.

**Table 1.** Participant characteristics.

| Characteristic | mvASD (n = 35) | TD (n = 30) |
| --- | --- | --- |
| Age (years) | 19.5 ± 4.2 (15–30) | 19.8 ± 4.0 (15–29) |
| Sex (F / M) | 9 / 26 | 10 / 20 |
| Mother tongue | Hebrew (100%) | Hebrew (100%) |
| Functional spoken words | < 30 (caregiver report) | Typical |
| SCQ total score, Completed by caregiver | 24.7 ± 6.1 | N/A |
| Diagnosis basis | DSM-IV / 5 criteria | N/A |
| Recruitment source | Clinics & special-ed schools | Community adverts |
| Key exclusions | Able to read aloud | Neurologic/psychiatric history |

### Stimuli and Procedures

#### Cued-Looking Paradigm (CLP)

We adapted the seminal Looking-While-Listening (LWL) paradigm (29), modifying it to present the cue word first and to use shorter stimulus durations. This adaptation allows testing of both text reading and speech comprehension. We refer to this as the “cued-looking” paradigm because the word— whether written or spoken—acts as a pre-cue or prime. The paradigm is illustrated in Figure 1.

**Figure 1.**
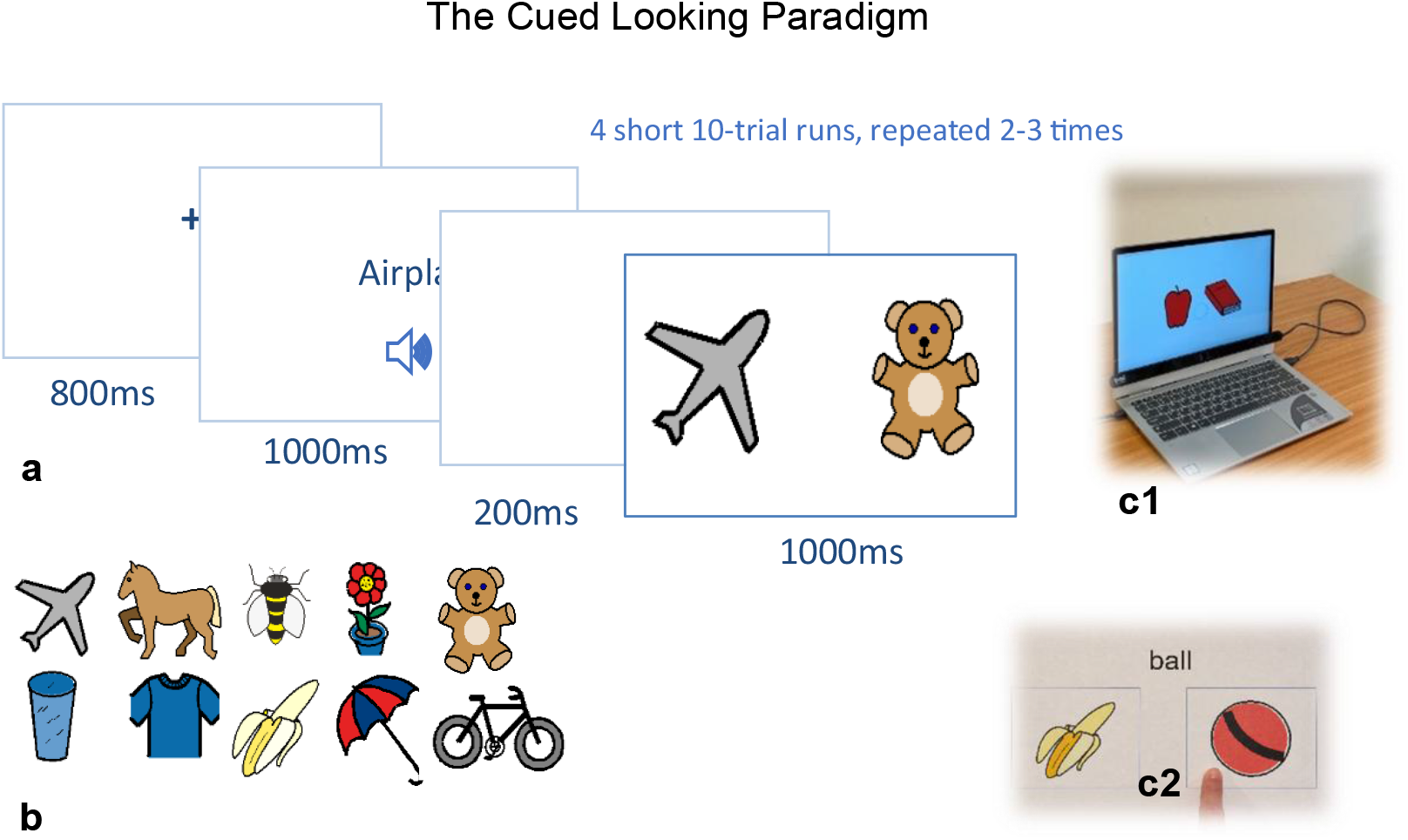
The Cued Looking Paradigm (CLP). (a) Trial sequence. Each trial begins with central fixation, followed by a cue (spoken or written word), a brief blank interval, and then two side-by-side pictures, target and foil. Display durations are indicated in the schematic. Participants are instructed to look at the named (spoken or written) item; gaze typically shifts automatically to the matching image. (b) Examples of the simple object drawings used as stimuli. (C1) Computer used to run the experiment, showing the task display on the screen. (c2) Example of a pointing-task card used in the experiment.

#### Stimuli

Color drawings of simple objects (examples in Figure 1b) were presented on a white background with luminance of ∼150 cd/m^2^. Images appeared ∼4 dva (degrees of visual angle) from fixation on the horizontal meridian (based on an average viewing distance of 54 cm) and typically subtended ∼4° in diameter. Their mean luminance was ∼70 cd/m^2^ (SD 25 cd/m^2^ for the inner parts), and the average RMS contrast across images was 0.472 (SD = 0.114), computed from pixel-wise relative luminance. In total, 28 images were used, taken from an educational image set (Widgit Symbols, The Center for Educational Technology, Tel Aviv, Israel). Text cues were presented in black Hebrew font (∼1 dva high and ∼3–5 dva wide). Spoken words were pre-recorded by a female colleague from the lab and played through the computer’s speaker at ∼67 dB SPL with a duration of ∼1 s (SD = 0.04 s).

#### Procedure

Experiments were run on a 13.3-inch Windows laptop with an FHD display at 60 Hz using the PSY experimental tool developed by author YSB, as in our previous studies (e.g., (30)). While observers fixated a central cross, a spoken or written cue was presented for 1 s, followed by a 200 ms blank screen and then a 1 s presentation of two side-by-side images (target and foil), one of which (the target) matched the cue (Figure 1a). Participants were instructed to look at the named object. Each experiment comprised four runs of 10 trials lasting 40–50 s each, repeated 2–3 times (∼100 trials per experiment). To sustain attention, a transparent “bubble” dynamically displayed real-time gaze position (a Tobii tracker feature). The experimenter (author KE) sat behind the participant, mostly invisible, to monitor tracking quality and avoid excessive redundant testing time. If tracking was interrupted, the run was repeated after a short or long break.

#### Pointing Paradigm

We conducted additional pointing experiments using the same vocabulary and images presented on printed cards. In the written cue condition, each standard A4 card (∼21 × 30 cm) displayed a text word and two images. Cards were shown until participants responded, and performance was recorded manually. Participants viewed the cards from ∼50 cm. The text spanned ∼5 × 1 cm, and each image measured 9 × 9 cm. A total of 40 cards were used, each presented once. The experiment was repeated with spoken words, in which the experimenter asked the participants to “Show me the object” referring to the target word. This format simulated conditions commonly encountered in educational settings and paper-based assessments used to evaluate cognitive abilities in individuals with severe autism.

### Eye-tracking and Data Analysis

Eye movements were recorded with a head-free eye tracker (Tobii 4C, 90 Hz); tracking precision was comparable across groups, as assessed by RMS sample-to-sample deviation (see SI Appendix). Horizontal gaze position was baseline-corrected and analyzed to derive two primary measures: (1) **gaze laterality** and (2) **eye-gaze performance**. Gaze laterality was defined as the difference in horizontal eye position between trials in which the target appeared on the right versus left, averaged over trials and aligned to picture onset. This measure captures the time course and magnitude of automatic gaze shifts toward the target image. **Eye-gaze performance** was computed on a per-trial basis by averaging horizontal gaze position within a predefined post-onset window (0.25–1.5 s) and scoring trials as correct when the sign (+ or-) of the averaged gaze position matched the target side (left matched negative). Trials with poor fixation during the cue period were excluded. Full preprocessing steps, calibration procedures, and robustness analyses are provided in the SI Appendix.

### Statistical analysis

Individual gaze laterality was assessed using time-resolved, non-parametric permutation tests comparing left- and right-target trials at the participant level (see SI.7 Appendix). Group comparisons were performed using two-tailed tests. Effect sizes (Cohen’s *d*) are reported where appropriate. Associations between behavioral and physiological measures were assessed using Pearson’s *r*, with Spearman rank correlations used to evaluate robustness to influential observations. Multiple comparisons across time points were controlled using false discovery rate (FDR) correction (*q* < 0.05). Uncertainty around regression estimates was visualized using bootstrap-derived confidence sleeves around orthogonal regression fits (see SI.6 Appendix). We quantified motor perseveration using two metrics computed from the sequence of responses in the pointing and eye-gaze tasks (see SI.5 Appendix).

## Results

### Gaze lateralization and performance in reading

Figure 2 presents the results of the Cued Looking Paradigm (CLP) for the reading condition. As described in Figure 1 and the Methods, gaze laterality was assessed by sorting horizontal eye-position traces by target side (left vs. right) and normalizing to the pre-stimulus baseline. Trials with unstable fixation during the reading period were excluded on a per-participant basis (see SI Appendix).

**Figure 2.**
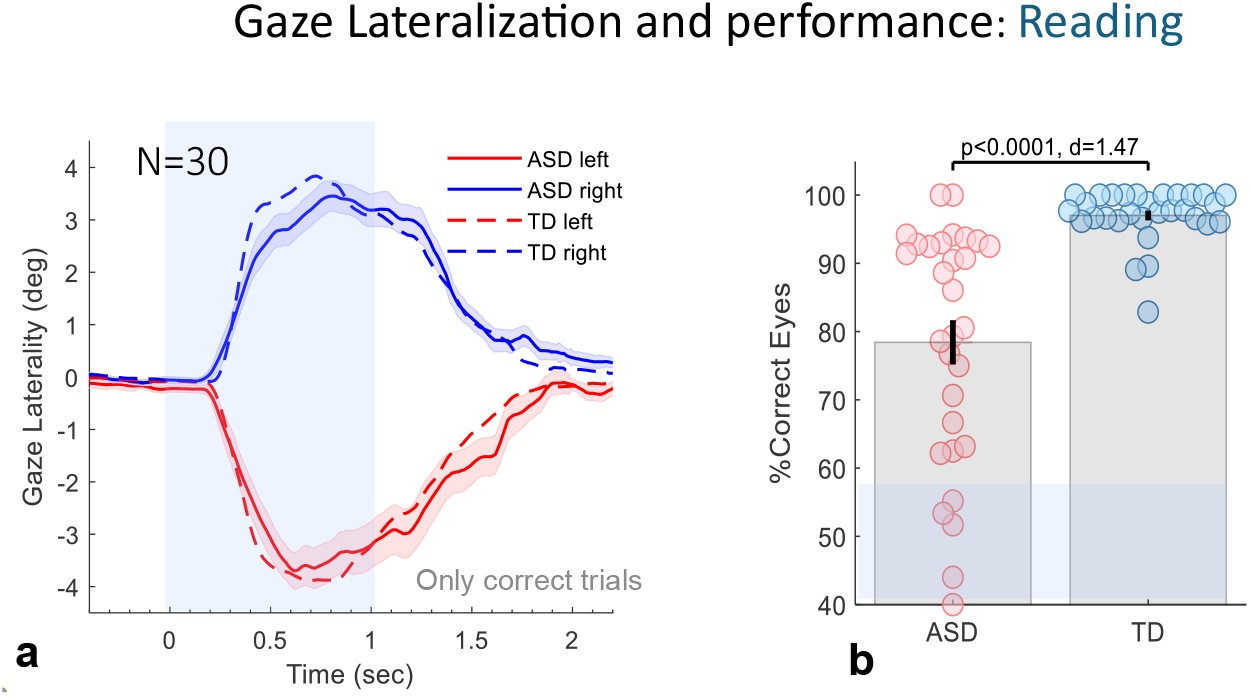
Gaze lateralization in the reading CLP. **(a)** Group-average horizontal eye-position traces for minimally verbal autistic (mvASD) participants and typically developing (TD) controls (dashed lines), including only trials with correct lateralization. Error sleeves denote ±1 SE across mvASD participants. Traces are sorted by target side (right: positive values in blue; left: negative in red). Gaze displacement toward the target image begins approximately 250 ms after image onset and follows a similar time course in both groups. **(b)** Eye-gaze performance (percent-correct lateralization) for the mvASD and TD groups shown as a bee-swarm plot. Each dot represents one participant, with group means ±1 SE overlaid. Significance (Wilcoxon rank-sum test) and Cohen’s *d* are shown. Shaded symbols denote participants with relatively low window-averaged gaze-laterality performance. Individual-level statistical significance was assessed separately using the time-resolved permutation analysis shown in Figure S1.

Individual gaze-laterality time courses for all 30 mvASD participants are shown in Figure S1. The majority of participants exhibited a clear lateralized gaze response toward the target image following picture onset. Using an individual-level, time-resolved nonparametric permutation test comparing left- and right-target trials (see SI.7), 24 of the 30 mvASD participants (80%) showed significant gaze lateralization in the expected direction, whereas 6 participants did not show reliable lateralization. Time intervals of significant lateralization for each participant are indicated by gray bars in Figure S1, based on the same permutation procedure.

Figure 2a shows group-average gaze traces for left- and right-target trials in the mvASD and TD groups, including only trials with correct lateralization and stable fixation. Both groups exhibited highly similar temporal dynamics and spatial amplitudes of gaze shifts, with gaze displacement toward the target image beginning approximately 250 ms after image onset. No group differences were observed in the timing of this gaze displacement. Likewise, maximal gaze laterality in the 0.25–1.5 s window was comparable between groups.

Figure 2b summarizes individual eye-gaze performance (percent-correct lateralization) for the reading CLP. While both groups performed well above chance, performance was lower on average in the mvASD group than in TD controls (78% vs. 97%). When excluding the six mvASD participants who did not show significant individual lateralization according to the time-resolved permutation analysis (Figure S1), the mvASD group average increased to 84.3%. Notably, a subgroup of 13 mvASD participants achieved performance levels above 90% correct, approaching the lower range of TD performance.

### Eye-gaze vs pointing in reading

In a separate experiment, all mvASD participants were tested on the same stimuli presented on laminated cards for pointing—each card containing the text word and two pictures (see Methods). The eye-gaze compared to pointing performance in the reading experiment are shown in Figures 3a, b. In striking contrast to their gaze-based reading, most mvASD participants who showed significant reading with their eyes performed at or near chance when pointing. This pattern is evident in Figure 3a, where most scatter-plot symbols lie above the diagonal (better performance with eyes than with pointing). Group-average performance is shown in Figure 3b, revealing a very low pointing performance (58%) and a highly significant difference between the two testing modes (t-test, p < 0.0001, Cohen’s d = 1.45). This effect was even stronger in the subgroup that scored above 80% with gaze: their average performance was 92% with the eyes versus 58% with pointing.

**Figure 3.**
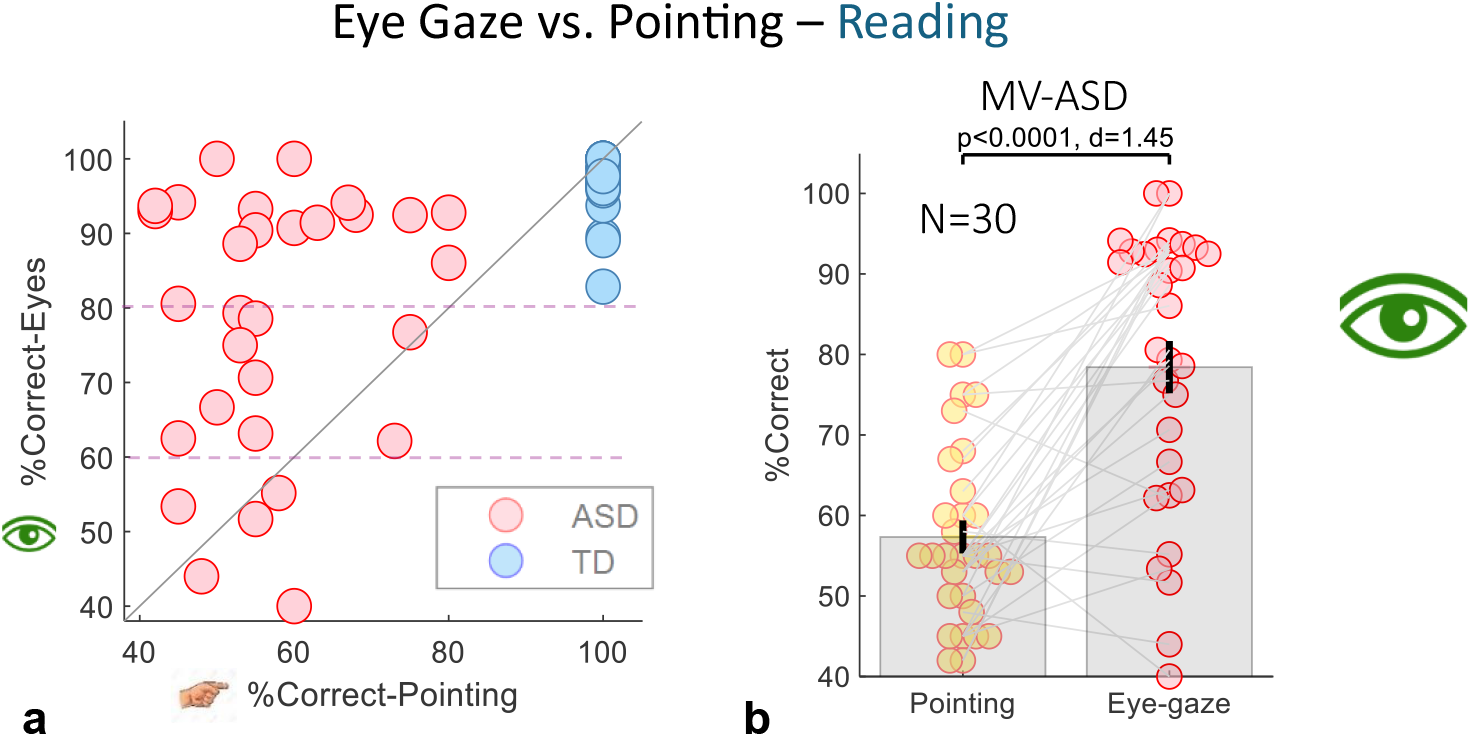
Eye-gaze versus pointing performance. (a,b) Eye-gaze and pointing performance in the reading experiment shown in two formats. (a) Scatter plot of pointing accuracy (x-axis) versus eye-gaze accuracy (y-axis), one dot per participant (red: mvASD; blue: TD controls). Most mvASD participants lie above the diagonal, indicating better performance with eye-gaze than with pointing. Sixteen mvASD participants scored above 80% correct, and thirteen above 90%. (b) Within-subject comparison of eye-gaze and pointing accuracy for reading. Lines connect data from the same participant; paired t-test p-values and Cohen’s d values are reported.

### Eye-gaze performance for hearing

In a second experiment, the text word was replaced with a spoken word (see Methods). Results are shown in Figure 4. Twenty-one of the 30 mvASD participants completed this experiment, which was administered after the reading condition and thus may have been affected by fatigue. Gaze performance in this experiment was slightly lower than in the reading condition, with the mvASD group averaging ∼78% (Figure 4a) compared to ∼82%, which was not significantly different (paired t-test, p=0.6). Importantly, there was no significant difference between pointing and eye-gaze in the listening experiment (Figure 4b, paired t-test, p=0.9).

**Figure 4.**
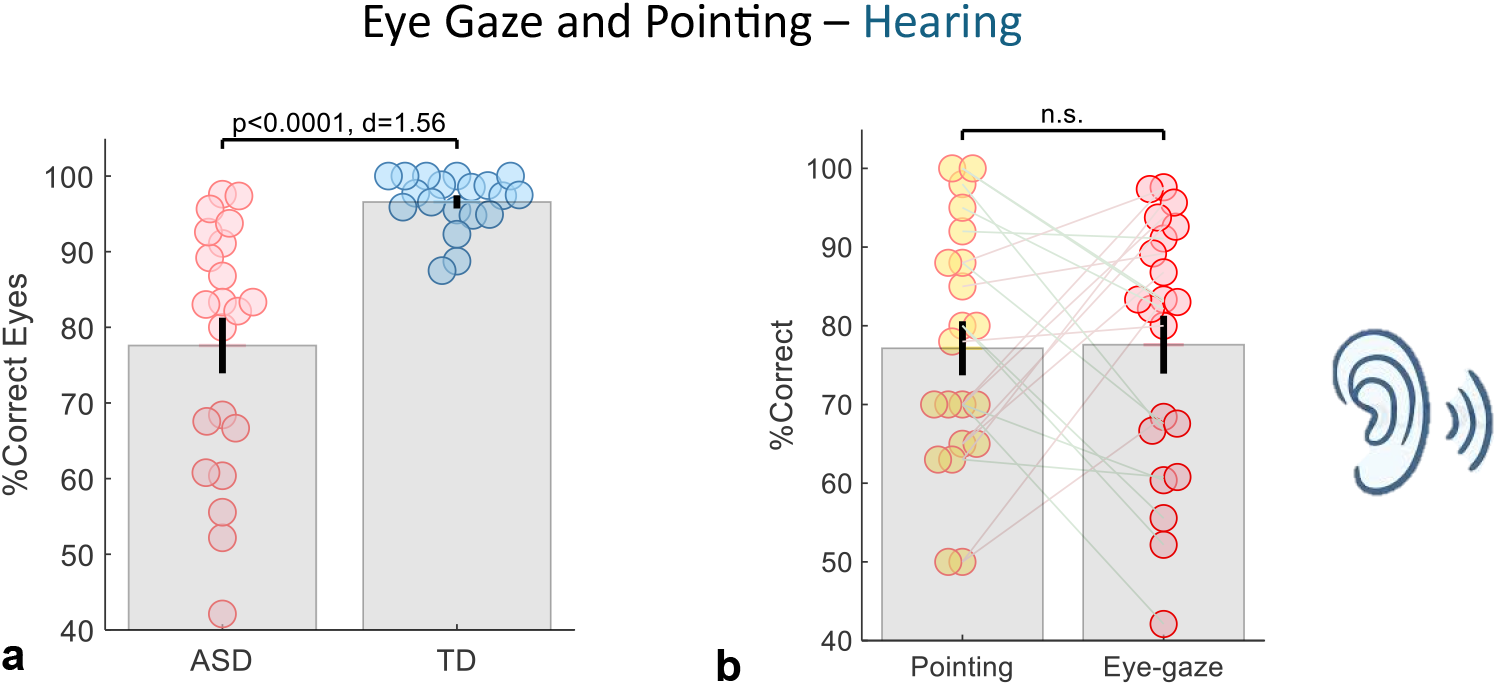
Results for the hearing experiment. (a) Percent-correct eye-gaze lateralization, with each dot representing one participant and group means ±1 SE overlaid. (b) Within-subject comparison of eye-gaze and pointing accuracy for hearing, analogous to Figure 3b.

### Perseveration measures

The two perseveration metrics revealed a clear dissociation between pointing and eye-gaze responses. For side bias, pointing responses showed significantly higher perseveration than gaze (p < 0.005, paired t-tests on both raw and z-normalized scores). The incongruent repetition index (IRI), a more behaviorally specific measure (Figure 5a), yielded the largest difference between modalities (p < 0.0001, d = 2.28), with similar results for the z-normalized scores (p < 0.0001, d = 3.48). Furthermore, the IRI showed a significant negative association with pupil dilation in the eye-gaze task (Figure 5b; Pearson r = –0.49, p < 0.007), indicating that greater perseveration in pointing was associated with reduced phasic arousal recruitment during gaze responses. This association was also present when assessed using Spearman rank correlation (p = 0.016) and remained significant after excluding one influential observation identified by a predefined ±3 SD orthogonal-regression residual criterion (Spearman p = 0.006). While the side bias measure produced some extreme values (e.g., z-scores > 8) in pointing responses, the incongruent repetition index provided a more balanced and interpretable distribution across participants and best captured the failure to adjust motor output in response to task-relevant changes.

**Figure 5.**
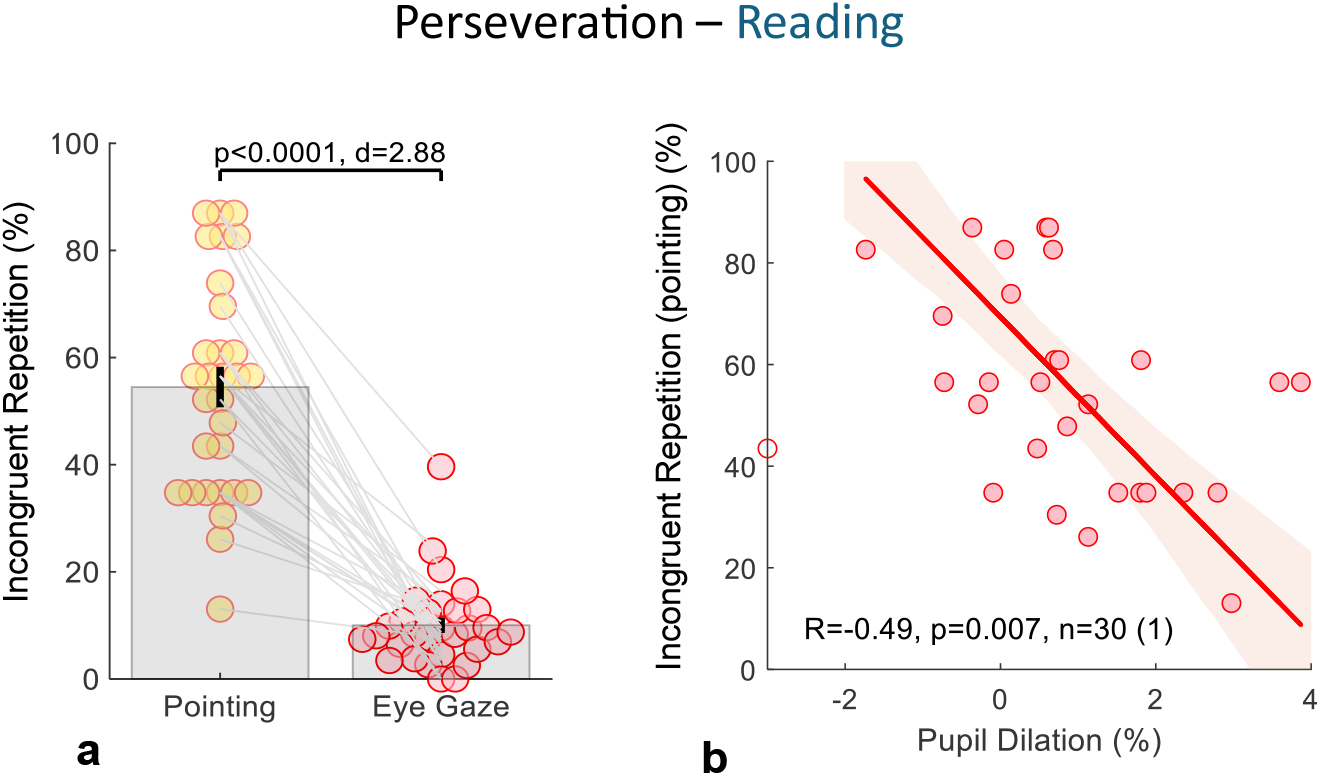
Perseveration in pointing responses (reading CLP). (a) Bee-swarm plot comparing perseveration rates—defined as responding to the same side when the target switches sides—between pointing and eye-gaze. The upper bar indicates t-test p-value and Cohen’s d. Pointing shows markedly higher perseveration than eye-gaze. (b) Association between pupil dilation during correct reading trials and pointing perseveration across participants. Greater pupil dilation is associated with less perseveration (Pearson *r* = –0.49, *p* = 0.007; *n* = 30 participants), computed after excluding one influential observation identified by a predefined ±3 SD orthogonal-regression residual criterion (thus, 29 data points contributed to the correlation). For visualization, the excluded observation is shown with an open symbol and plotted at – 3 on the y-axis (true value –4.8). Shaded regions denote 95% bootstrap confidence sleeves around the orthogonal regression fit (see SI.6 for details).

### Pupil dilation and gaze performance

Pupil dilation in response to the pictures may reflect both changes in luminance and the recruitment of arousal needed to make a decision and shift gaze. We analyzed pupil-size changes for the two groups in the reading experiment, with results shown in Figure 6. First, we compared the time course of pupil change (percentage relative to the pre-onset baseline; see Methods) in both experiments. As shown in Figure 6a, the TD group exhibited much stronger dilation, peaking around 0.5 s after picture presentation. We quantified dilation per trial by computing the maximum in the 0.5–1 s window after stimulus onset; individual and group averages (including only trials classified as correct based on gaze laterality; see Methods) are shown in Figure 6b. The TD group showed markedly larger dilation than the mvASD group (5.5% vs. 1.3%, p < 0.0005). This difference in dilation could not have been due to differences in exposure to the images since in this analysis only correct trials were taken into account, and the average gaze lateralization for correct trials was very similar between groups (Figure 2b). Figure 6c shows the correlation between pupil dilation and eye-gaze performance (correct trials only). A significant positive relationship emerged between the change from baseline (in %) and eye-gaze performance. The correlation shown in Figure 6c was significant using both Pearson and Spearman correlations (Spearman p = 0.0022), indicating a robust association not driven by extreme values.

**Figure 6.**
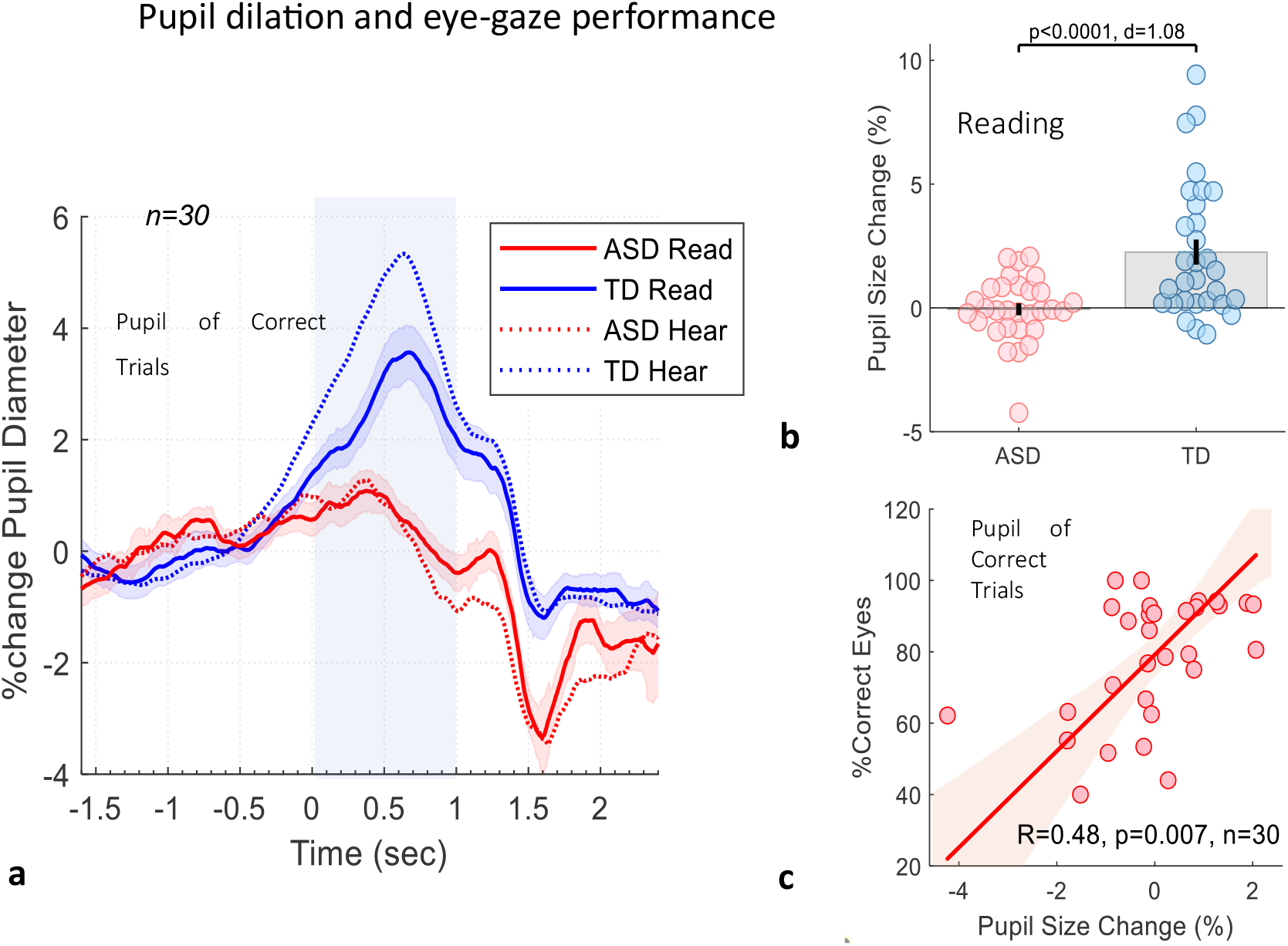
Pupil size change at picture onset in the CLP. (a) Time course of pupil size change (percentage relative to the pre-onset baseline) averaged across participants. Solid lines: reading; dotted lines: hearing. Shaded areas show ±1 SE. (b) Mean pupil change 0–1 s post-onset for correct trials in reading, plotted as group means ±1 SE with individual data points overlaid. mvASD participants show markedly reduced dilation compared with TD controls (n = 30 each). (c) Correlation between pupil size change from (b) and eye-gaze performance for each mvASD participant. No participants exceeded the predefined outlier criterion (±3 SD of orthogonal-regression residuals); therefore, no data points were excluded. Shaded regions denote 95% bootstrap confidence sleeves around the orthogonal regression fit (see SI.6 for details).

## Discussion

### Revealing Hidden Abilities via Eye Gaze: Summary of Findings

The Cued Looking Paradigm (CLP), which presents spoken/written words *prior* to visual stimuli measuring eye-gaze laterality, offers, to our knowledge the first systematic direct evidence of hidden reading and receptive language skills in individuals with mvASD. These findings challenge assumptions of severe cognitive and language impairments and extend recent reports of strengths in communication and cognition found in some individuals (3). A clear dissociation between gaze-based performance and traditional pointing responses suggests that standard assessment methods may significantly underestimate cognitive abilities in this population.

### Gaze–Pointing Dissociation and the Role of Automaticity

Most mvASD adolescents and adults showed a clear reading effect during the CLP, similar to typical development (TD) controls. This was evident in their consistent lateral gaze toward target image. In the written-word condition, mvASD participants demonstrated similar reaction times and saccade amplitudes to TD participants. Similar results were found in the spoken-word condition. However, many of the same individuals performed at or near chance when asked to point to the correct image on laminated cards. This striking dissociation could reflect the fact that, unlike eye gaze— a highly overlearned and largely automatic response (16) —pointing remains an effortful, non-automatized skill for many minimally verbal individuals, making it a challenge in testing situations. Similar gaze–pointing dissociations have been observed by our group in other tasks, including a simple visual oddball paradigm with older children (Sykes-Haas & Bonneh, preprint (10)) supporting the view that these effects generalize beyond the present CLP and are not task-specific artifacts.

### CLP Paradigm Strengths vs. Prior Work

Our findings differ substantially from earlier studies that used eye tracking to assess receptive language in mvASD. For example, Plesa-Skwerer et al. (19) reported only ∼60% accuracy in the Looking-While-Listening (LWL) paradigm, with better performance when participants pointed (72%). In our CLP, by contrast (Figure 3), mvASD participants achieved 87% accuracy in the spoken-word condition and 79% in the written-word condition, while performing significantly worse when pointing (58% in reading). CLP results also diverge from ERP studies, which often detect group-level effects but show limited reliability at the individual level (18). In the CLP data, robust gaze-based effects were evident for most participants, indicating the paradigm provides a more sensitive and individually valid measure of hidden lexical-semantic abilities. We propose the structure of the CLP is central to its effectiveness. Unlike traditional paradigms like LWL, which present auditory and visual information simultaneously, the CLP presents the spoken or written cue before the test images. This sequencing allows the cue to act as a prime, reducing visual competition and facilitating rapid, automatic orienting to the target.

In addition, the CLP presents information sequentially and unimodally—one stimulus type at a time—thereby reducing the cognitive demands of integrating input across sensory channels that are often atypical in ASD (31). This likely lowers task load and helps individuals with suspected LC–NE dysregulation sustain engagement (32) consistent with evidence for atypical arousal regulation in mvASD (see below). The CLP’s temporal structure is also analogous to “gap” trials in oculomotor research, where removing a central fixation before target onset shortens saccadic latency by reducing competition between stimuli (33, 34). In contrast, LWL and similar paradigms resemble “overlap” trials, in which simultaneous stimulus presentation increases competition and slows orienting—a challenge that may be particularly pronounced in autism.

Taken together, these design features—sequential presentation, reduced competition, and unimodal presentation—may explain why the CLP elicits clearer and more consistent results than prior methods. The remaining question is *why* mvASD participants succeed with gaze but struggle with deliberate motor responses, such as pointing, an issue addressed below.

### Pupil Dilation and LC-NE Dysregulation

To explore a potential mechanism accounting for the gap between comprehension and expression in mvASD, we examined pupil dilation (PD) during CLP trials. PD is considered an indirect index of activity in the locus coeruleus–norepinephrine (LC-NE) system, which plays a key role in attention and the initiation of voluntary behavior (32). Compared to neurotypical controls, mvASD participants showed reduced pupil dilation, especially those with lower gaze-based performance (Figure 6b). This was also reflected in a significant correlation between pupil size and eye-gaze task accuracy (Figure 6c), and in an inverse correlation with pointing perseveration (Figure 5b). Our results support the view that LC-NE dysregulation links atypical arousal to autistic cognition and difficulties in choosing and initiating actions (32), potentially through reduced transient recruitment of arousal required for deliberate actions such as pointing. In contrast, more automatic behaviors such as gaze shifts may remain accessible, offering a powerful alternative for assessing latent cognitive abilities. The CLP’s continuous, computerized presentation with fixed and predictable timing likely places lower demands on transient arousal recruitment than clinician-led pointing tasks, which require repeated re-engagement and adjustment to variable timing, voice, and stimulus placement.

### Dysregulated Action Selection as a Mechanism for Hidden Cognition

The convergence of pupil dilation, perseveration, and head-movement findings points to a dysregulation of the neural systems that govern action selection. Within the *affordance-competition* framework (14), multiple potential actions—such as gaze shifts or pointing movements—compete for execution based on sensory evidence. In minimally verbal autism, this competition may be imbalanced: voluntary, goal-directed actions like pointing fail to be initiated or stabilized, whereas highly automatized actions such as gaze shifts remain accessible. Reduced phasic activation of the locus coeruleus– norepinephrine (LC-NE) system, reflected in diminished pupil dilation, may weaken the transient arousal signal needed to resolve this competition in favor of the intended action. The CLP design effectively bypasses this bottleneck by minimizing competing motor and sensory demands, allowing latent skills to manifest through largely automatic eye movements rather than effortful motor output.

### Limitations

Our findings should be interpreted in light of several limitations. The vocabulary tested consisted of simple, concrete nouns typically acquired in early childhood, and we did not examine sentence comprehension, grammar, or phonological decoding. Thus, our results do not imply high-level language or general cognitive competence. Sample bias is also a concern: many participants were recruited through networks of parents who believe their children have unrecognized abilities, although we also observed similar patterns among 7 participants from a special-education setting. The examiner (KE) was not blinded to study aims; while unintentional cueing is unlikely given the rapid timing of eye movements, and the mostly out-of-view position of the examiner, future studies should consider blinded examiners for eye-gaze and pointing conditions. Sessions were lengthy (40–90 minutes) and required a skilled professional to interpret subtle behaviors and maintain cooperation, which may limit scalability. Finally, participants pointed to laminated cards rather than a touchscreen. Pointing is often inconsistently acquired and not automatized in mvASD (5, 17), and previous touchscreen studies have likewise not shown good pointing performance on somewhat similar tasks (19). We therefore expect that the core dissociation between pointing and gaze would persist if our pointing tasks were assessed on a touch screen.

### Implications and Future Directions

Our findings indicate that some minimally verbal individuals with autism possess lexical–semantic abilities that remain hidden in traditional assessments. By removing the need for intentional motor responses, the Cued Looking Paradigm (CLP) offers a new way to access these abilities. Future research could extend CLP to more complex tasks—such as simple sentence comprehension (35), verb and action-word processing, and phonological awareness (e.g., presenting spoken nonwords paired with visually similar written forms) to determine to what extent minimally verbal individuals rely on whole-word recognition versus phoneme–grapheme decoding, a distinction relevant to hyperlexia and atypical reading development (36). Beyond language, CLP may also be adapted to nonverbal cognition such as categorization, memory, or visual reasoning. Furthermore, early identification of hyperlexic abilities could help tailor instruction for young minimally verbal children who can already recognize written words.

## ACKNOWLEDGMENTS

This work was supported by a grant from the Simons Foundation Autism Research Initiative (SFARI; Grant No. 573840), and by the Israel Science Foundation Grant 657/21 (both to YSB.).

## SI Appendix

### SI Methods

#### SI.1 Eye-tracking hardware and calibration

Participants’ eye movements were recorded with a Tobii 4C eye tracker sampling at 90 Hz, which we found to be reliable and robust for head-free tracking (see our recent pupil paper (1), our preprint on pursuit and saccades in TBI (2) and others (3)). This device was chosen for its reliability and flexibility in working with the severely autistic. Accuracy was verified in preliminary experiments with controls, yielding a typical fixation SD of ∼0.5 dva in the current setup. This precision is also evident in the main results, including effects of pupil-size changes of <1%. Before each session, the tracker was calibrated using Tobii’s fast four-point calibration. Some mvASD participants required several attempts. In a few cases, calibration failed; for these participants we used calibration data from another participant with a similar profile (see below for how calibration inaccuracy was handled).

#### SI.2 Eye-tracking data quality

To quantify tracking precision, we computed the root-mean-square sample-to-sample deviation (RMS S2S) of gaze position during fixation periods, using the two-dimensional (horizontal and vertical) gaze signal. RMS S2S was calculated as the square root of the mean squared Euclidean displacement between successive samples, excluding missing data. RMS S2S values did not differ significantly between mvASD and TD participants (mvASD: mean ± SD = 0.081 ± 0.029; TD: mean ± SD = 0.087 ± 0.044 dva; *p* = 0.59), indicating comparable tracking precision across groups and sufficient data quality to resolve the multi-degree gaze displacements analyzed in the CLP.

#### SI.2 Eye-tracking preprocessing

Because participants viewed the stimuli with unrestricted head movements (see Procedure), actual viewing distance varied across trials and participants. For each trial we estimated viewing distance as the mean eye-to-screen distance from –0.4 to 0 s relative to tracking onset, excluding >2 SD outliers. Average participant distances ranged from 42–74 cm (one control outlier at a higher distance). Group means were similar (controls 56.5 cm; mvASD 54.1 cm; p = 0.28). To standardize gaze data across participants for group-level averaging, eye positions were converted from screen pixels to degrees of visual angle using a fixed reference distance of 54 cm. Data were preprocessed to detect and remove periods without tracking and blinks (including preceding vertical displacements) with a 50 ms margin, following our previous studies (4). Epochs were extracted from –2 s to +3 s relative to picture onset (Figure 1a). Participants with fewer than 20 valid epochs (epochs containing >50% valid samples) were excluded, leaving 30 mvASD participants. Additional epochs were rejected if fixation during the reading period (–1.2 to –0.2 s before image presentation) was inaccurate (Figure 1a). To mitigate calibration inaccuracy, we computed each participant’s baseline fixation during the reading period (averaged across runs) and marked as invalid any epoch whose displacement from baseline exceeded a threshold. A 4 dva threshold was used for testing individual significance, and a stricter 2 dva threshold for performance analyses (rejecting more trials but improving accuracy). On average, this procedure led to the exclusion of 45% of trials in the mvASD group and 15% in the TD group, reflecting greater fixation instability in mvASD participants. In general, outliers exceeding 2 SD from the mean were removed from all computed averages.

#### SI.3 Gaze laterality and performance computation

Two primary measures were derived from the CLP: (1) Gaze Laterality (Figures 2a,b) and (2) Eye-gaze Performance (Figure 2c). *Gaze laterality* was computed by averaging horizontal gaze traces of valid epochs sorted by target side (left or right), rejecting >2 SD outliers at each time point, and baseline-correcting (subtracting the mean gaze position from –2 to –0.2 s before image presentation). This analysis was used to visualize and quantify the time course of lateralized gaze, which then defined the time window for the performance measure. *Eye-gaze performance* (percent correct) was computed by averaging the horizontal gaze position per trial in the 0.25–1.5 s window after image onset (again removing >2 SD outliers and baseline-correcting using the –2 to 0 s period, termed “Hav”). A trial was scored as correct when the sign of Hav matched the side of the target. Invalid trials (deviant fixation during reading; 2 dva threshold) were excluded. For each participant, we assessed above-chance laterality by testing the difference between Hav values for left-versus right-sided targets using one-tailed t-tests and Wilcoxon rank-sum tests with FDR correction (5) across participants, excluding trials with deviant fixation (>4 dva). Group comparisons used two-tailed t-tests, and we additionally computed effect sizes (Cohen’s d).

#### SI.4 Pupil preprocessing and normalization

Pupil analysis: Pupil dilation was analyzed by first averaging left-eye pupil traces from valid epochs (good fixation; see above) with correct lateralization, rejecting outliers (>2 SD from the mean) at each time point and converting the values to a percentage relative to baseline (the epoch’s average pupil size during the last 1 s before image presentation). This procedure was used for visualization and for determining the region for peak detection. We then computed the maxima and minima of the pupil traces, as well as the mean percentage change from baseline (1 s pre-image period) in the 0.5–1 s window after image presentation.

#### SI.5 Perseveration analysis

We quantified motor perseveration using two metrics computed from the sequence of responses in the pointing and eye-gaze tasks. The first and simplest measure was *side bias*, defined as the absolute deviation from equal left/right responses: 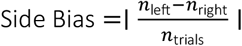. This captures the extent to which a participant preferentially pointed to one side across trials. In addition, we computed an *incongruent repetition index*, designed to reflect motor perseveration in the face of task demands. Specifically, we measured how often a participant repeated the same response (left or right) despite a change in the correct side from the previous trial. The metric was calculated as the proportion of trials where the response repeated but the correct side had changed:

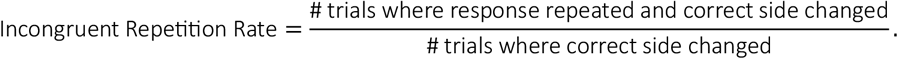

A high value indicates a tendency to perseverate in response despite shifting task contingencies. To assess whether incongruent repetition rates exceeded chance expectations, we generated null distributions by simulating 1000 random response sequences per subject, matched in length and trial structure to the observed data. Each subject’s observed incongruent repetition index was compared to this distribution to compute a Z-score and empirical one-tailed p-value.

#### SI.6 Bootstrap procedures for confidence sleeves

Confidence sleeves around regression lines were computed using nonparametric bootstrap resampling. From the original set of paired observations (*x*_*i*_, *y*_*i*_) (after excluding NaNs), we generated 1,000 bootstrap datasets by sampling *N*pairs with replacement (sample size equal to the original *N*). For each bootstrap dataset, an orthogonal regression line was fit by applying PCA to the z-scored (*x, y*) values and taking the first principal axis as the best-fitting direction; the resulting slope was then transformed back to the original measurement units and the intercept computed from the sample means. Each fitted line was evaluated on a fixed grid of 100 x-values spanning the observed x-range with a ±5% margin. At each grid point, the 2.5th and 97.5th percentiles of the bootstrap-predicted y-values defined the 95% confidence sleeve. These sleeves represent uncertainty around the estimated regression relationship (confidence band for the fitted line), not prediction intervals for individual data points.

#### SI.7 nonparametric permutation test

To determine time intervals showing reliable gaze lateralization, we applied a nonparametric permutation-based test (6, 7) comparing left- and right-target trials. At each time point, the difference in horizontal gaze position between the two conditions was first computed across participants. To account for temporal dependence and avoid parametric assumptions, statistical significance was evaluated using a permutation procedure. Specifically, target-side labels were randomly reassigned within participants, and the group-level gaze difference time course was recomputed for each of 1,000 permutations. For each permutation, the maximal test statistic across time was extracted to form a null distribution. Observed time points exceeding the 95th percentile of this distribution were considered significant. This approach controls for multiple comparisons across time while preserving the temporal structure of the data.

#### SI.8 Robustness analyses

Individual gaze lateralization was assessed using two complementary approaches. First, time-resolved differences between left- and right-target conditions were evaluated within each participant using a non-parametric permutation test (Figure S1). Second, gaze lateralization was quantified as the mean horizontal eye displacement within a predefined post-stimulus time window (main text), yielding a scalar measure per participant. Both approaches identified a largely overlapping subset of participants showing reliable lateralization in the expected direction, indicating that individual-level effects were not dependent on a specific statistical test or temporal aggregation choice. While the two approaches yielded highly consistent results, a minor discrepancy was observed in the number of participants classified as showing reliable lateralization. The time-resolved permutation analysis (Fig. S1) identified significant lateralization in 24 of 30 participants (80%), whereas the window-averaged gaze lateralization measure (Figure 2b) classified 25 of 30 participants (83%) as showing correct lateralization. This difference reflects the stricter temporal requirements of the permutation-based analysis. For reporting purposes, we adopt the more conservative estimate (80%), noting that both analyses provide convergent evidence for robust individual-level gaze lateralization.

Robustness of the correlation-based findings was evaluated using alternative correlation metrics. For the association shown in Figure 4b, one participant exhibited an extreme pupil value that affected the Pearson correlation; however, the relationship remained significant when assessed using Spearman rank correlation (p = 0.006). For Fig. 5c, no extreme values were present, and the association was significant using both Pearson and Spearman correlations (Spearman p = 0.0022). These analyses indicate that the reported associations reflect stable monotonic relationships rather than being driven by a small number of influential observations.

### SI Figures

**Figure S1.**
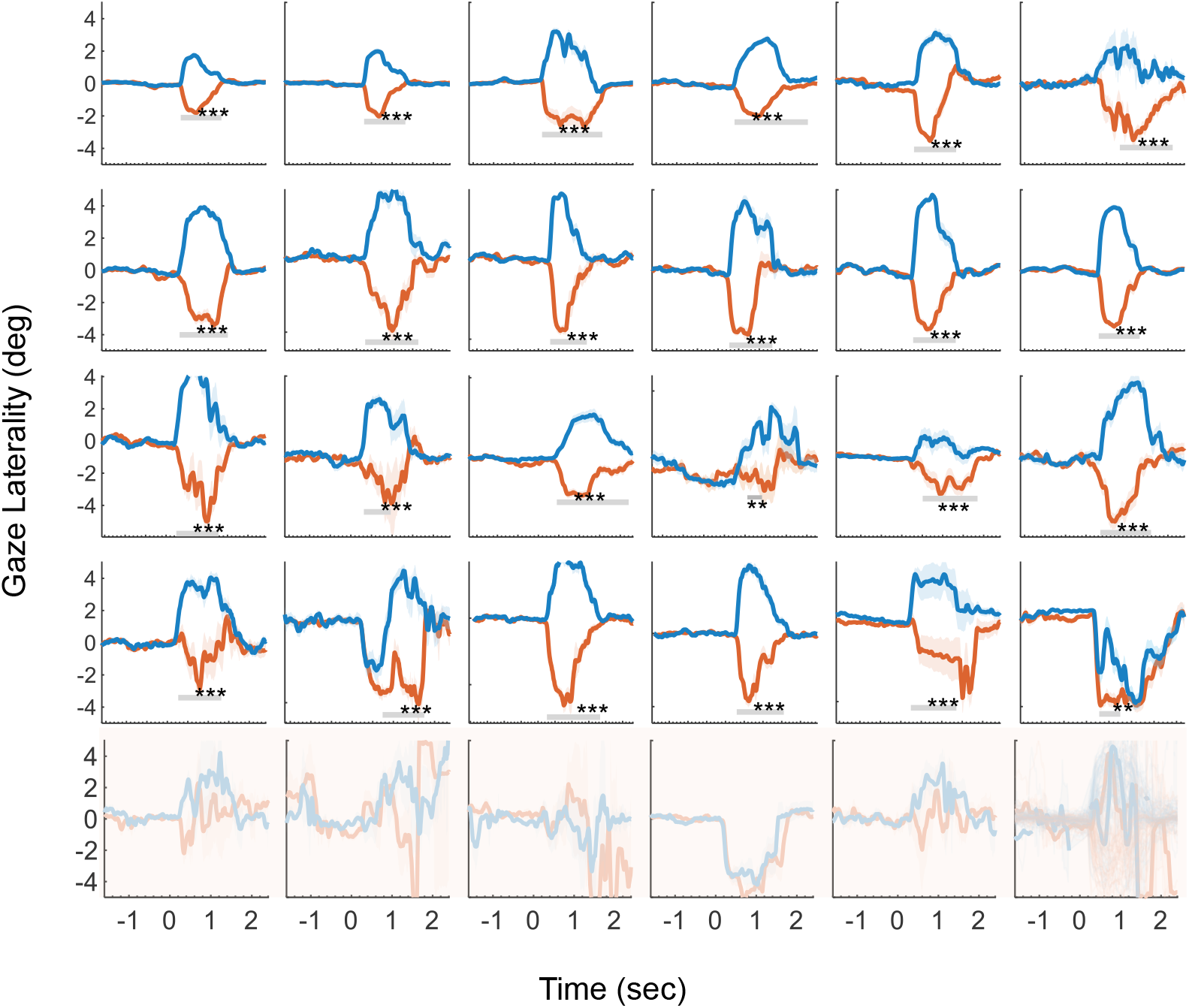
Individual gaze lateralization during the reading CLP task. Horizontal gaze position over time is shown for all 30 minimally verbal autistic (mvASD) participants, averaged across trials (correct and incorrect). Shaded sleeves denote ±1 SEM across trials within each participant. Traces are color-coded by target side (right: blue, positive values; left: red, negative values). The top four rows show 24 participants who exhibited significant gaze lateralization in the expected direction, whereas the bottom row shows 6 participants who did not show reliable lateralization at the individual level. Gray horizontal bars indicate time intervals in which gaze position differed significantly between left and right target conditions, assessed using a non-parametric permutation test (see Methods).

## References

1. B. Chakrabarti, Commentary: Critical considerations for studying low-functioning autism. J. Child Psychol. Psychiatry 58, 436–438 (2017).

2. H. Tager-Flusberg, C. Kasari, Minimally verbal school-aged children with autism spectrum disorder: The neglected end of the spectrum. Autism Res. 6, 468–478 (2013).

3. M. Pizzano, et al., Profiles of minimally verbal autistic children: Illuminating the neglected end of the spectrum. Autism Res. 17, 1218–1229 (2024).

4. C. Kasari, N. Brady, C. Lord, H. Tager-Flusberg, Assessing the minimally verbal school-aged child with autism spectrum disorder. Autism Res. 6, 479–493 (2013).

5. V. Courchesne, A. A. S. Meilleur, M. P. Poulin-Lord, M. Dawson, I. Soulières, Autistic children at risk of being underestimated: School-based pilot study of a strength-informed assessment. Mol. Autism 6, 4–13 (2015).

6. Y. Chen, B. Siles, H. Tager-Flusberg, Receptive language and receptive-expressive discrepancy in minimally verbal autistic children and adolescents. Autism Res. 17, 381–394 (2024).

7. R. S. Eagle, Accessing and Assessing Intelligence in Individuals with Lower Functioning Autism. (2002).

8. V. H. Bal, C. Farmer, A. Thurm, Describing Function in ASD: Using the DSM-5 and Other Methods to Improve Precision. J. Autism Dev. Disord. 47, 2938–2941 (2017).

9. K. D. Bello, N. Goharpey, S. G. Crewther, D. P. Crewther, A puzzle form of a non-verbal intelligence test gives significantly higher performance measures in children with severe intellectual disability. BMC Pediatr. 8, 30 (2008).

10. Y. S. B. Hannah S. Sykes-Haas, Minimally verbal children with autism may ‘see the point, but do not (always) point to what they see.’ [Preprint] (2025).

11. K. Boser, S. Higgins, A. Fetherston, M. A. Preissler, B. Gordon, Semantic Fields in Low-Functioning Autism.

12. N. Bauminger-Zviely, et al., Communicating Without Words: School-Based RCT Social Intervention in Minimally Verbal Peer Dyads with ASD. J. Clin. Child Adolesc. Psychol. 49, 837–853 (2020).

13. M. Kedar, N. Bauminger-Zviely, Predictors of individual differences in minimally verbal peer communication exchanges following PEER-ORIENTED social intervention. Autism Res. 16, 230–244 (2023).

14. P. Cisek, Cortical mechanisms of action selection: The affordance competition hypothesis. Philos. Trans. R. Soc. B Biol. Sci. 362, 1585–1599 (2007).

15. E. B. Torres, et al., Autism: The Micro-Movement perspective. Front. Integr. Neurosci. 7, 1–26 (2013).

16. R. K. Mishra, C. N. L. Olivers, F. Huettig, Spoken language and the decision to move the eyes: To what extent are language-mediated eye movements automatic? Prog. Brain Res. 202, 135–149 (2013).

17. M. A. Gernsbacher, E. A. Sauer, H. M. Geye, E. K. Schweigert, H. Hill Goldsmith, Infant and toddler oral- and manual-motor skills predict later speech fluency in autism. J. Child Psychol. Psychiatry 49, 43–50 (2008).

18. C. DiStefano, D. Senturk, S. S. Jeste, ERP evidence of semantic processing in children with ASD. Dev. Cogn. Neurosci. 36, 100640 (2019).

19. D. Plesa-Skwerer, S. E. Jordan, B. H. Brukilacchio, H. Tager-Flusberg, Comparing methods for assessing receptive language skills in minimally verbal children and adolescents with autism spectrum disorders. Autism 20, 591–604 (2016).

20. D. Orlievsky, S. Cukier, Language, writing, and activity disorder in the autistic spectrum. Front. Integr. Neurosci. 7, 1–3 (2013).

21. M. P. Mostert, Facilitated Communication since 1995: A Review of Published Studies. J. Autism Dev. Disord. 31, 287–313 (2001).

22. Y. S. Bonneh, et al., When they see, they see it almost right: Normal subjective experience of detected stimuli in spatial neglect. Neurosci. Lett. 446, 51–55 (2008).

23. I. Kedar, Ido in Autismland: Climbing Out of Autism’s Silent Prison (2012).

24. V. K. Jaswal, A. J. Lampi, K. M. Stockwell, Literacy in nonspeaking autistic people. Autism 28, 2503–2514 (2024).

25. V. K. Jaswal, A. Wayne, H. Golino, Eye-tracking reveals agency in assisted autistic communication. Sci. Rep. 10, 7882 (2020).

26. L. Wing, A. Shah, A Systematic Examination of Catatonia-like Clinical Pictures in Autism Spectrum Disorders. Int. Rev. Neurobiol. 72, 21–39 (2006).

27. C. Cantiani, et al., From sensory perception to lexical-semantic processing: An ERP study in non-verbal children with autism. PLoS ONE 11, 1–31 (2016).

28. E. L. Coderre, et al., Implicit Measures of Receptive Vocabulary Knowledge in Individuals with Level 3 Autism. Cogn. Behav. Neurol. 32, 95–119 (2019).

29. A. Fernald, R. Zangl, Looking while listening: Using eye movements to monitor spoken language. Dev. Psycholinguist. 97–135 (2008).

30. Y. S. Bonneh, Y. Adini, U. Polat, Contrast sensitivity revealed by spontaneous eyeblinks: Evidence for a common mechanism of oculomotor inhibition. J. Vis. 16 (2016).

31. A. B. Brandwein, et al., The Development of Multisensory Integration in High-Functioning Autism: High-Density Electrical Mapping and Psychophysical Measures Reveal Impairments in the Processing of Audiovisual Inputs. Cereb. Cortex 23, 1329–1341 (2013).

32. E. B. London, Neuromodulation and a Reconceptualization of Autism Spectrum Disorders: Using the Locus Coeruleus Functioning as an Exemplar. Front. Neurol. 9, 1–16 (2018).

33. M. Elsabbagh, et al., Disengagement of visual attention in infancy is associated with emerging autism in toddlerhood. Biol. Psychiatry 74, 189–194 (2013).

34. B. Keehn, G. Kadlaskar, S. Bergmann, R. McNally Keehn, A. Francis, Attentional Disengagement and the Locus Coeruleus – Norepinephrine System in Children With Autism Spectrum Disorder. Front. Integr. Neurosci. 15 (2021).

35. F. Huettig, J. Audring, R. Jackendoff, A parallel architecture perspective on pre-activation and prediction in language processing. Cognition 224, 105050 (2022).

36. A. Ostrolenk, B. Forgeot d’Arc, P. Jelenic, F. Samson, L. Mottron, Hyperlexia: Systematic review, neurocognitive modelling, and outcome. Neurosci. Biobehav. Rev. 79, 134–149 (2017).

## References

1. R. Meidan, Y. S. Bonneh, Pattern-induced visual discomfort and its cumulative effects revealed by pupillary measures. Front. Hum. Neurosci. 19, 1723675 (2026).

2. S. Shani, Bedside Assessment of Visual Tracking in Traumatic Brain Injury: Comparing Simple and Predictive Paradigms Using Multiple Oculomotor Markers.

3. T. Lee, M. Yeung, S. Sze, A. Chan, Eye-Tracking Training Improves Inhibitory Control in Children with Attention-Deficit/Hyperactivity Disorder. Brain Sci. 11, 314 (2021).

4. Y. S. Bonneh, Y. Adini, U. Polat, Contrast sensitivity revealed by spontaneous eyeblinks: Evidence for a common mechanism of oculomotor inhibition. J. Vis. 16 (2016).

5. Y. Benjamini, Y. Hochberg, Controlling the False Discovery Rate: A Practical and Powerful Approach to Multiple Testing. J. R. Stat. Soc. Ser. B Methodol. 57, 289–300 (1995).

6. A. Widmann, R. Engbert, E. Schroger, Microsaccadic Responses Indicate Fast Categorization of Sounds: A Novel Approach to Study Auditory Cognition. J. Neurosci. 34, 11152–11158 (2014).

7. E. Maris, R. Oostenveld, Nonparametric statistical testing of EEG- and MEG-data. J. Neurosci. Methods 164, 177–190 (2007).

